# Collapse Precedes Folding in Denaturant-Dependent Assembly of Ubiquitin

**DOI:** 10.1101/081299

**Authors:** Govardhan Reddy, D. Thirumalai

## Abstract

Despite the small size the folding of Ubiquitin (Ub), which plays an indispensable role in targeting proteins for degradation and DNA damage response, is complex. A number of experiments on Ub folding have reached differing conclusions regarding the relation between collapse and folding, and whether intermediates are populated. In order to resolve these vexing issues, we elucidate the denaturant-dependent thermodynamics and kinetics of Ub folding in low and neutral pH as a function of Guanidinium chloride and Urea using coarse-grained molecular simulations. The changes in the fraction of the folded Ub, and the radius of gyration (*R*_*g*_) as a function of the denaturant concentration, [*C*], are in quantitative agreement with experiments. Under conditions used in experiments, *R*_*g*_ of the unfolded state at neutral pH changes only by ≈ 17% as the [*GdmCl*] decreases from 6 M to 0 M. We predict that the extent of compaction of the unfolded state increases as temperature decreases. A two-dimensional folding landscape as a function of *R*_*g*_ and a measure of similarity to the folded state reveals unambiguously that the native state assembly is preceded by collapse, as discovered in fast mixing experiments on several proteins. Analyses of the folding trajectories, under mildly denaturing conditions ([*GdmCl*]=1.0M or [*Urea*]=1.0M), shows that Ub folds by collision between preformed secondary structural elements involving kinetic intermediates that are primarily stabilized by long-range contacts. Our work explains the results of Small Angle X-Ray Scattering (SAXS) experiments on Ub quantitatively, and establishes that evolved globular proteins are poised to collapse. In the process, we explain the discrepancy between SAXS and single molecule fluorescent resonant energy transfer (smFRET) experiments, which have arrived at a contradicting conclusion concerning the collapse of polypeptide chains.

## Introduction

Mono and poly-ubiquitin play an important role in cell signaling pathways. Ubiquitination of specific lysine residues in the target proteins^1^ is a signal for triggering cellular processes such as protein degradation by the proteosomes,^2^ and DNA damage response needed for genome stability.^3^ Thus, understanding how Ubiquitin (Ub) folds is important in describing its cellular functions. The crystal structure^4^ (PDB ID: 1UBQ) of the 76 residue monomeric Ub shows that the folded state has five *β*-sheets and two helices (Figure 1A). The *C*_*α*_ contact-map based on *α*-carbon atom shows that there are short-range contacts between the amino acid residues in the helices (*α*_1_ and *α*_2_), and the hairpin *β*_1_*β*_2_. Long range contacts connect the strands *β*_1_-*β*_5_, *β*_3_-*β*_5_, and the loops *L*_1_-*L*_2_ (Figure S1). The predicted complex Ub folding kinetics^5^ as a function of temperature is attributed to the multiple long-range contacts in the folded state of Ub setting the stage for an in depth investigation of how chemical denaturants modulate the folding landscape of Ub.

**Figure 1:**
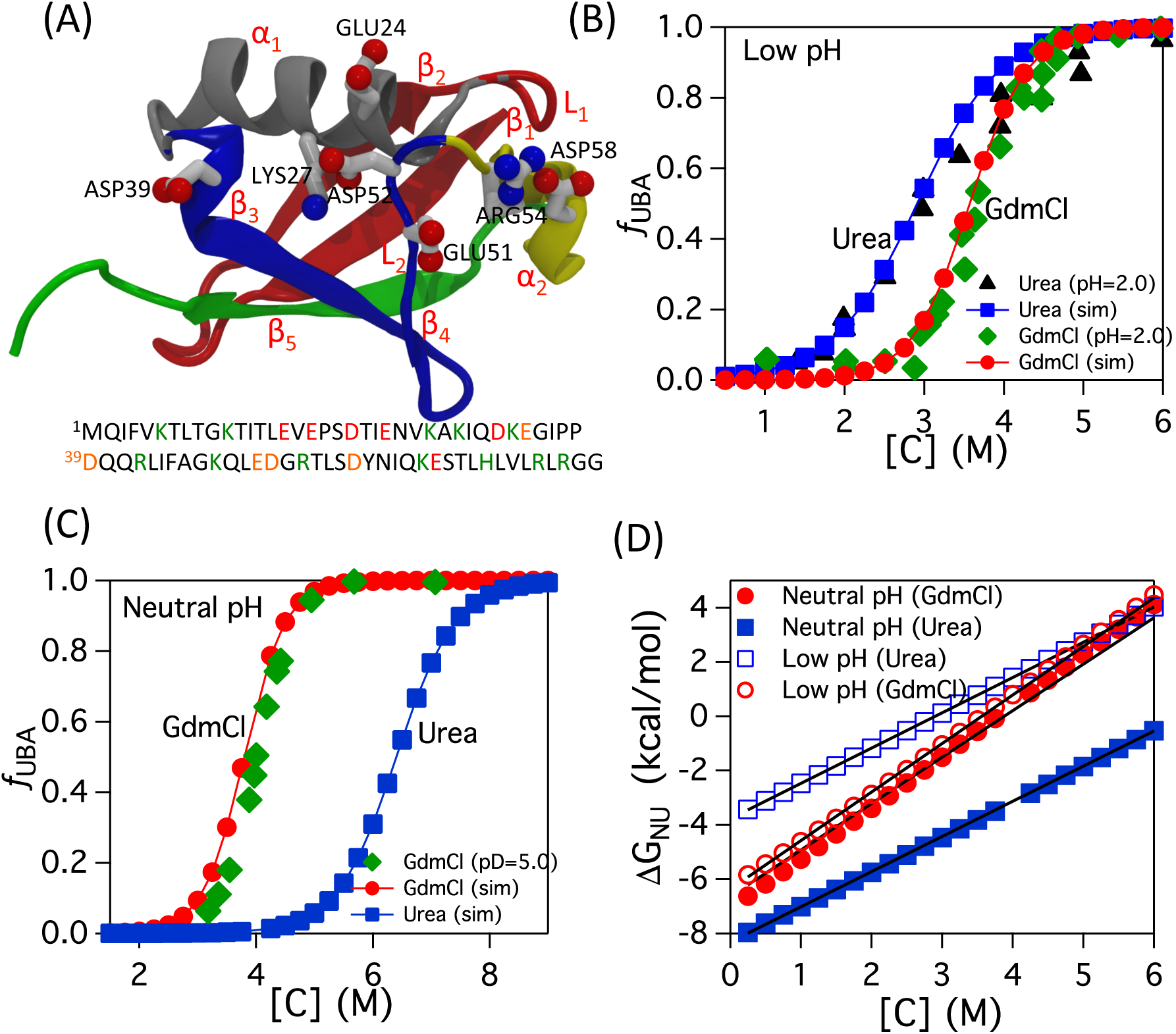
Denaturant-dependent folding of Ub. (A) Crystal structure of Ub (PDB ID: 1UBQ). The folded structure consists of 5 *β*-sheets labeled *β*_1_ (red), *β*_2_ (red), *β*_3_ (blue), *β*_4_ (blue) and *β*_5_ (green). Two helices are shown in *α*_1_ (silver) and *α*_2_ (yellow). Charged residues at the interface of the helix *α*_1_ and loops *L*_1_ and *L*_2_ are shown. At the bottom the single letter codes for the amino acid residues present in Ub is shown. The letters in green and red are positively and negatively charged amino acids, respectively. (B) The fraction of the protein in the UBA, *f*_*UBA*_ as a function of the denaturant concentration [*C*] in low pH. The data in solid green diamonds and solid black triangles is *f*_*UBA*_ as function of [*GdmCl*] and [*Urea*] respectively from experiments^39^ performed at pH = 2.0. The data in solid red circles and blue squares is [*GdmCl*] and [*Urea*] data respectively from simulations using the low pH Ub model. (C) The data in solid green diamonds is *f*_*UBA*_ as a function of [*GdmCl*] from experiments^36^ performed at pD = 5.0. Data in solid red circles and blue circles is [*GdmCl*] and [*Urea*] data respectively from simulations using the neutral pH Ub model. (D) Free energy difference between the folded and unfolded states, ∆*G*_*NU*_ as a function of [*C*].

A combination of experiments^6–9^ and theory^10–15^ has yielded insights into the generic mechanisms by which proteins fold. However, it has been difficult to provide a molecular description of folding mechanisms, which are experimentally probed using denaturants whereas most of the folding simulations are done by varying temperature. Because of the large differences in the impact of temperature and denaturants on the various states of proteins direct comparisons between experiments and computations is largely elusive. This is a major bottleneck in assessing the efficacy of simulations. A major advance in overcoming this bottleneck was made with the introduction of the phenomenological Molecular Transfer Model (MTM)^16,17^ that combines the classical transfer model and coarse-grained representation of the polypeptide chain. In several previous studies^16–21^ MTM has been applied to *quantitatively* describe denaturant-dependent folding of a number of proteins containing between 50-250 amino acid residues.

Here, we investigate the thermodynamics and kinetics of Ub folding, which has been investigated by a variety of experiments^22–39^ and computations.^5,40–54^ However, the effects of denaturants such as Guanidinium chloride and Urea, have not been considered in simulations. This is important because there are few central experimental controversies, such as the extent of collapse in the denatured ensemble of Ub as the denaturant concentration is lowered and if intermediates are present, that remain unresolved. We provide quantitative insights into these problems by dissecting Ub folding using simulations accounting for both denaturant and pH effects.

Ub has a significant number of charged residues, making the folding properties dependent on pH.^5,38,39,55^ By incorporating electrostatic interactions within the framework of the SOP-SC model^19,56^ we performed exhaustive simulations over a wide range of temperature and denaturant concentrations. Various thermodynamic properties of Ub computed from simulations, such as the fraction of the protein in the unfolded basin of attraction *f*_*UBA*_, the free energy difference between the folded and unfolded state ∆*G*_*NU*_, and the radius of gyration *R*_*g*_, as a function of the denaturant concentrations are in excellent agreement with experiments.^36,39,57^ The pair distance distribution functions *P*(*r*), and *R*_*g*_ computed for the burst phase of Ub folding show that the size of Ub decreases only modestly in the early stages of folding, in agreement with the Small Angle X-ray scattering experiments (SAXS).^57^ The equilibrium values of the radius of gyration 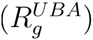 of the unfolded ensemble decreases continuously as the denaturant concentration is lowered. Folding landscape as a function of *R*_*g*_ and the structural overlap function (*χ*) shows that compaction precedes folding. Ub in highly stabilizing conditions folds through the diffusion-collision model by populating distinct kinetic intermediates with partially folded structures. Our study tidily resolves the apparent controversies between conclusions reached using SAXS and smFRET experiments, and suggests that the tendency to collapse is universal of all evolved globular proteins.

## Methods

### Self Organized Polymer-Side Chain (SOP-SC) model

As described in detail else-where,^5^ we modeled Ub using a coarse-grained SOP-SC model.^19,56^ Each residue is represented using two interaction centers, one for the backbone atoms and the other for the side chain (SC). The interaction centers are at the *C*_*α*_ atom position of the residue, and the center of mass of the side chain. The SCs interact via a residue-dependent statistical potential.^58^ Acidic residues (Figure 1A) are protonated at low pH, minimizing the effect of electrostatic interactions. To mimic Ub folding in neutral pH, we added charges by placing them on the side chains of the charged residues. The SOP-SC models for Ub are constructed using the crystal structure.^4^

The force field in the SOP-SC model is a sum of bonded and non-bonded interactions. The bonded interactions (*E*_*B*_), between a pair of connected beads (two successive *C*_*α*_ atoms or a SC connected to a *C*_*α*_ atom), account for chain connectivity. The non-bonded interactions are a sum of native 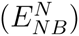 and non-native 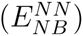 interactions. If two beads are separated by at least three bonds, and if the distance between them in the coarse-grained crystal structure is less than a cutoff distance *R*_*c*_ (Table S1) then their interactions are considered native. The rest of the pairs of beads, not covalently linked or native, are classified as non-native interactions. Electrostatic effects at neutral pH are modeled using the screened Coulomb potential (*E*^*el*^).^5^

The coarse-grained force-field of a protein conformation in the SOP-SC model represented by the coordinates **{r}** at [*C*] = 0 is given by

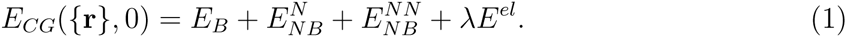

Description of the various energy terms in Equation 1 can be found in the SI. The parameter *λ* can take the values 0 or 1 to either switch off or switch on the electrostatic effects. The parameters used in the energy function are in Table S1 in the SI.

### Molecular Transfer Model

We used the Molecular Transfer Model (MTM) model^16,17^ to compute the Ub folding thermodynamics and kinetics in the presence of denaturants, Urea and Guanidinium Hydrochloride. In a solution at denaturant concentration [*C*], the effective coarse-grained force field for the protein using MTM framework is,

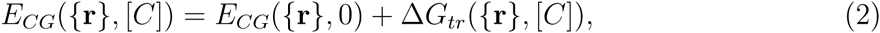

where *E*_*CG*_(**{r}**, 0) is given by Equation 1. The protein-denaturant interaction free energy (∆*G*_*tr*_(**{r}**, [*C*])) in an aqueous solution at a denaturant concentration [C] is given by

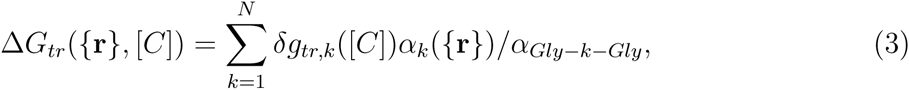

where *N*(=*N*_*res*_ × 2 = 152) is the number of residues in Ub, *δg_tr,k_*([*C*]) is the transfer free energy of bead *k*, *α_k_*(**{r}**) is the solvent accessible surface area (SASA) of the site *k* in a protein conformation described by positions **{r}**, *α*_*Gly*−k−*Gly*_ is the SASA of the bead *k* in the tripeptide *Gly–k–Gly*. The radii for side chains of amino acids needed to compute *α_k_*(**{r}**) are given in Table S2 in Ref.^19^ The experimental^16,18,59^ transfer free energies *δg_tr,i_*([*C*]), which depend on the chemical nature of the denaturant, for backbone and side chains are listed in Table S3 in Ref.^19^ The values for *α*_*Gly*−k−*Gly*_ are listed in Table S4 in Ref.^19^

### Simulations

We used low friction Langevin dynamics simulations^60^ to obtain the thermodynamic properties. The average value of a physical quantity, *A*, at any temperature, *T* and [C] is calculated^16,17^ using the Weighted Histogram Method (WHAM),^61^

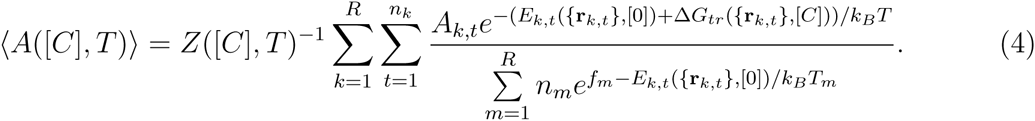

In Equation 4, *R* is the number of simulation trajectories, *n*_*k*_ is the number of protein conformations from the *k*^*th*^ simulation, *A*_*k,t*_ is the value of the property of the *t*^*th*^ conformation from the *k*^*th*^ simulation, *T*_*m*_ and *f*_*m*_ are the temperature and free energy respectively from the *m*^*th*^ simulation, *E_k,t_*(**{r**_*k,t*_}, [0]) and ∆*G*_*tr*_(**{r**_*k,t*_}, [*C*])) are the internal energy at [*C*] = 0 M and MTM energy respectively of the *t*^*th*^ conformation from the *k*^*th*^ simulation. The partition function *Z*([*C*]*, T*) is,

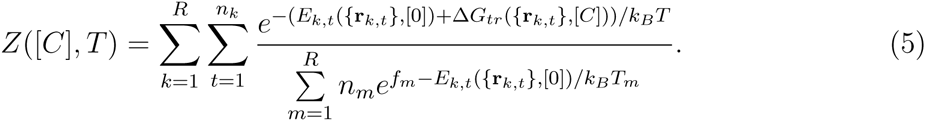

We performed Brownian dynamics simulations^62^ to simulate the protein folding kinetics. To obtain the folding trajectories at the concentration [*C*], the full Hamiltonian (Equation 2) with a friction coefficient corresponding to water (see SI for details) is used.

### Data Analysis

The structural overlap function,^63^ 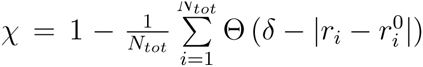, distinguishes the naive basin of attraction (NBA) and unfolded basin of attraction (UBA); *N*_*tot*_(= 11026) is the number of pairs of interaction centers in the SOP-SC model of Ub separated by at least 2 bonds, *r*_*i*_ is the distance between the *i*^*th*^ pair of beads, and 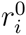 is the distance in the folded state, Θ is the Heaviside step function, and *δ* = 2 Å. A plot of *χ* as a function of time, and the probability distribution of *χ* at the melting temperature *T*_*m*_, in both neutral and acidic pH are shown in Figure S2 in the SI. The fraction of molecules in the unfolded basin of attraction (UBA) as a function of the denaturant concentration, [*C*], is calculated using *χ* as an order parameter (see SI for details). The radius of gyration is calculated using 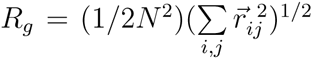, where 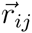 is the vector connecting interaction centers *i* and *j*.

In order to determine the order of formation of contacts that are separated along the sequence, we used the local contact order,^64,65^ *S*_*i*_. We define *S*_*i*_ associated with residue *i* as,

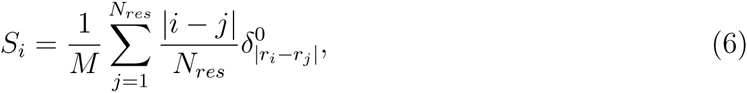

where *|i − j|* is the length of the protein sequence between the residues *i* and *j*, *M* is the number of native contacts that the *i*^*th*^ residue forms with the other residues in the folded state, 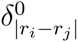 is the Kronecker delta function used to count the native contacts formed by the residue *i*, and superscript 0 indicates that only native contacts are considered.

## Results

### Thermodynamics of Ubiquitin folding

We computed the fraction of Ub in the unfolded basin of attraction (*f*_*UBA*_), radius of gyration (*R*_*g*_), and the free energy difference between the native and unfolded state (∆*G*_*NU*_ = *G_NBA_ − G_UBA_*) as a function of the denaturant concentration ([*C*]) in both low pH and neutral pH (Figure 1).

### Linear extrapolation method is valid in Urea

In order to compute the thermodynamic properties as a function of [*Urea*], we choose *T* = 343.25 *K*, since ∆*G*_*NU*_ (*T* = 343.25 *K,* [*C*] = 0 *M*) *≈* 3.23 *kcal/mol* is approximately equal to the experimental value ∆*G*_*NU*_ (*T* = 298 *K,* [*Urea*] = 0 *M*) at pH = 2.0.^39^ We choose *T* = 333 *K* to calculate the [*GdmCl*]-dependent properties, since ∆*G*_*NU*_ (*T* = 333 *K,* [*C*] = 0 *M*) *≈* 6.6 *kcal/mol* is approximately equal to the ∆*G*_*NU*_ (*T* = 298 *K,* [*GdmCl*] = 0 *M*) found in experiments.^39^ The simulation temperatures in the two denaturants are different because experiments^39^ at pH = 2.0 show that the ∆*G*_*NU*_ ([*C*] = 0 *M*) values extracted using Urea and GdmCl experiments do not coincide. The value ∆*G*_*NU*_ ([*Urea*] = 0 *M*) = 3.23 *kcal/mol* obtained from Urea denaturation experiments agrees with the differential scanning calorimetry experiments^66^ in water indicating that the linear extrapolation method (LEM) used to extrapolate ∆*G*_*NU*_ ([*Urea*]) as function of [*Urea*] data to obtain ∆*G*_*NU*_ ([*Urea*] = 0 *M*), is reliable. The LEM method fails for [*GdmCl*] experiments, as the plot of ∆*G*_*NU*_ ([*GdmCl*]) as a function of [*GdmCl*] for concentrations [*GdmCl*] *<* 1.0 *M* is not linear. When this plot is linearly extrapolated to [*GdmCl*] = 0 *M*, it yields ∆*G*_*NU*_ ([*GdmCl*] = 0 *M*) = 6.6 *kcal/mol*, approximately twice the value obtained in differential scanning experiments.^66^

To compute thermodynamic properties as a function of denaturant concentration in neutral pH we choose *T* = 333.25 *K* ensuring that ∆*G*_*NU*_ (*T* = 333.25 *K,* [*C*] = 0 *M*) *≈* 7.5 *kcal/mol* is approximately equal to ∆*G*_*NU*_ (*T* = 298 *K,* [*GdmCl*] = 0 *M*) from experiments^36^ at pD = 5.0. Experiments for Ub denaturation in neutral pH by Urea are not available. We used this procedure of choosing simulation temperatures to compute properties in different denaturing conditions because we cannot get absolute free energies using CG models. It is worth emphasizing that this is the only adjustable parameter in the MTM model.

### Population of the unfolded state as a function of [*C*]

The calculated *f*_*UBA*_ in low and neutral pH as a function of [*GdmCl*] is in excellent agreement with experiments^36,39^ (Figure 1B and 1C) as are the predictions for Urea denaturation in low pH^39^ (Figure 1B). It is interesting that in low pH, Urea acts as a better denaturing agent than GdmCl. The denaturing ability of the Guanidinium ions depends on the anion^67^ as well as pH, which modulates the electrostatic interactions. In neutral pH the large concentrations for both Urea and GdmCl required to denature Ub reflects the extraordinary stability of Ub. The free energy difference between the folded and unfolded states, ∆*G*_*NU*_, as a function of [*C*] in both low and neutral pH is linear in both GdmCl and Urea (Figure 1D). The slopes of the lines, representing the *m* values, in low pH for GdmCl and Urea are *≈* 1.8 *kcal mol*^−1^ *M*^−1^ and 1.3 *kcal mol*^−1^ *M*^−1^, respectively. The *m*-values from experiments^39^ in low pH for GdmCl and Urea are 1.8 and 1.0 *kcal mol*^−1^ *M*^−1^ respectively. The computed *m*-values for GdmCl and Urea in neutral pH are *≈* 1.7 *kcal mol*^−1^ *M*^−1^ and 1.3 *kcal mol*^−1^ *M*^−1^, respectively. The *m*-value for GdmCl denaturation computed from simulations compares well with the experimental^36^ value of 1.9 *kcal mol*^−1^ *M*^−1^.

### Radius of gyration as a function of [*C*]

The equilibrium radius of gyration, 〈*R*_*g*_〉, as a function of [*GdmCl*] from simulations is in excellent agreement with SAXS measurements^57^ at neutral pH (Figure 2A). Our simulations also reproduce the midpoint value of the denaturant concentration ([*GdmCl*] *≈* 3.8 M) where the protein folding-unfolding transition occurs in neutral pH (Figure 2A). Experimental data for *R*_*g*_ as a function of [*Urea*] or [*GdmCl*] in low pH is not available for comparison with the simulation data (Figure 2B). The 〈*R*_*g*_〉 of the unfolded ensemble depends on pH. Experiments^57,68,69^ estimate that *R*_*g*_ of the unfolded state at pH 2.5 is *≈* 32 Å whereas it is more compact at pH 7.0 with *R_g_ ≈* 26 Å. In neutral pH, electrostatic interactions play a dominant role and contribute to the compaction of the protein.^5^ In low pH, the acidic residues are protonated minimizing the role of electrostatic interactions and the protein samples conformations with higher *R*_*g*_ values.^5^

**Figure 2:**
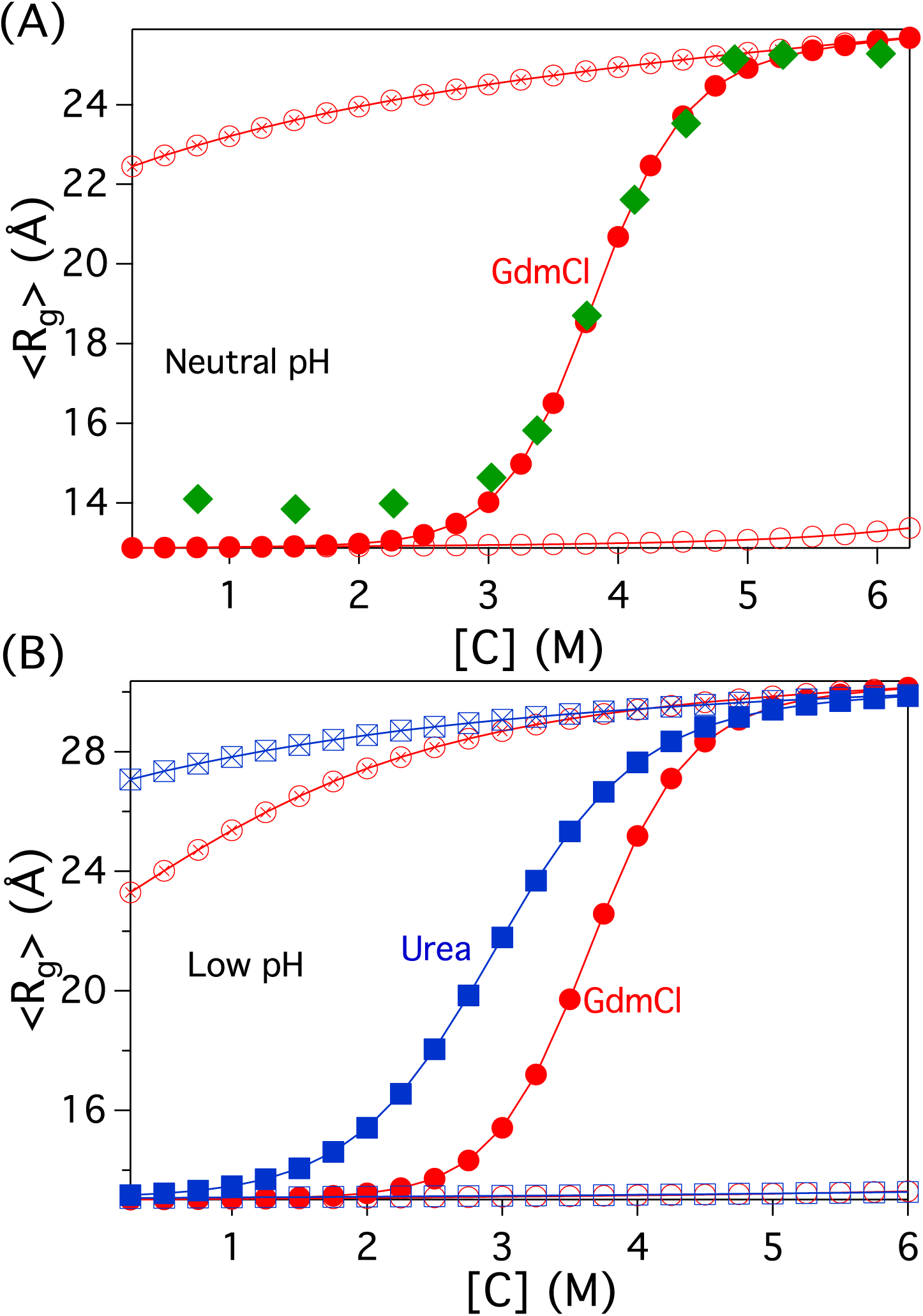
The radius of gyration, *R*_*g*_, as a function of denaturant concentration, [*C*]. (A) Neutral pH. Data in red solid circles and green diamonds are from simulations and experiments, ^57^ respectively for [*GdmCl*]. 〈*R*_*g*_〉 of UBA and NBA basins computed from simulations are shown in red circles with cross and red empty circles, respectively. (B) Low pH. Symbols in red circles and blue squares are for [*GdmCl*] and [*Urea*], respectively. Empty symbols and symbols with cross represent NBA and UBA basins, respectively.

### Unfolded states are compact under native conditions

Despite considerable evidence to the contrary there is doubt, based largely on SAXS experiments on protein L and Ub, that the radius of gyration of the UBA, 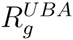 remains a constant even at low [*C*]. The excellent agreement between simulations and experiments in Figure 2A allows us to shed light on the puzzling conclusions reached elsewhere.^57^ To this end we calculated 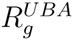 using only the ensemble of unfolded conformations. The results in Figure 3 unambiguously show that 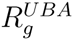, decreases in both low and neutral pH as [*GdmCl*] decreases from 6 M to 0 M indicating that the protein on an average decreases in size. At pH = 7.0, 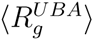 decreases from *≈* 26 Å to *≈* 22 Å, where as in low pH, *R*_*g*_ decreases from *≈* 30 Å to *≈* 23 Å (Figure 2). In low pH, on dilution of [*Urea*] from 6 M to 0 M, the change in *R*_*g*_ is only *≈* 3 Å, which is less than the *≈* 7 Å decrease upon [*GdmCl*] dilution. A lower temperature (*T* = 333 *K*) is used [*GdmCl*] simulations (*T* = 333 *K*) compared to [*Urea*] (*T* = 343.25 *K*) accounting (in part) for the larger compaction as [*GdmCl*] is decreased.

**Figure 3:**
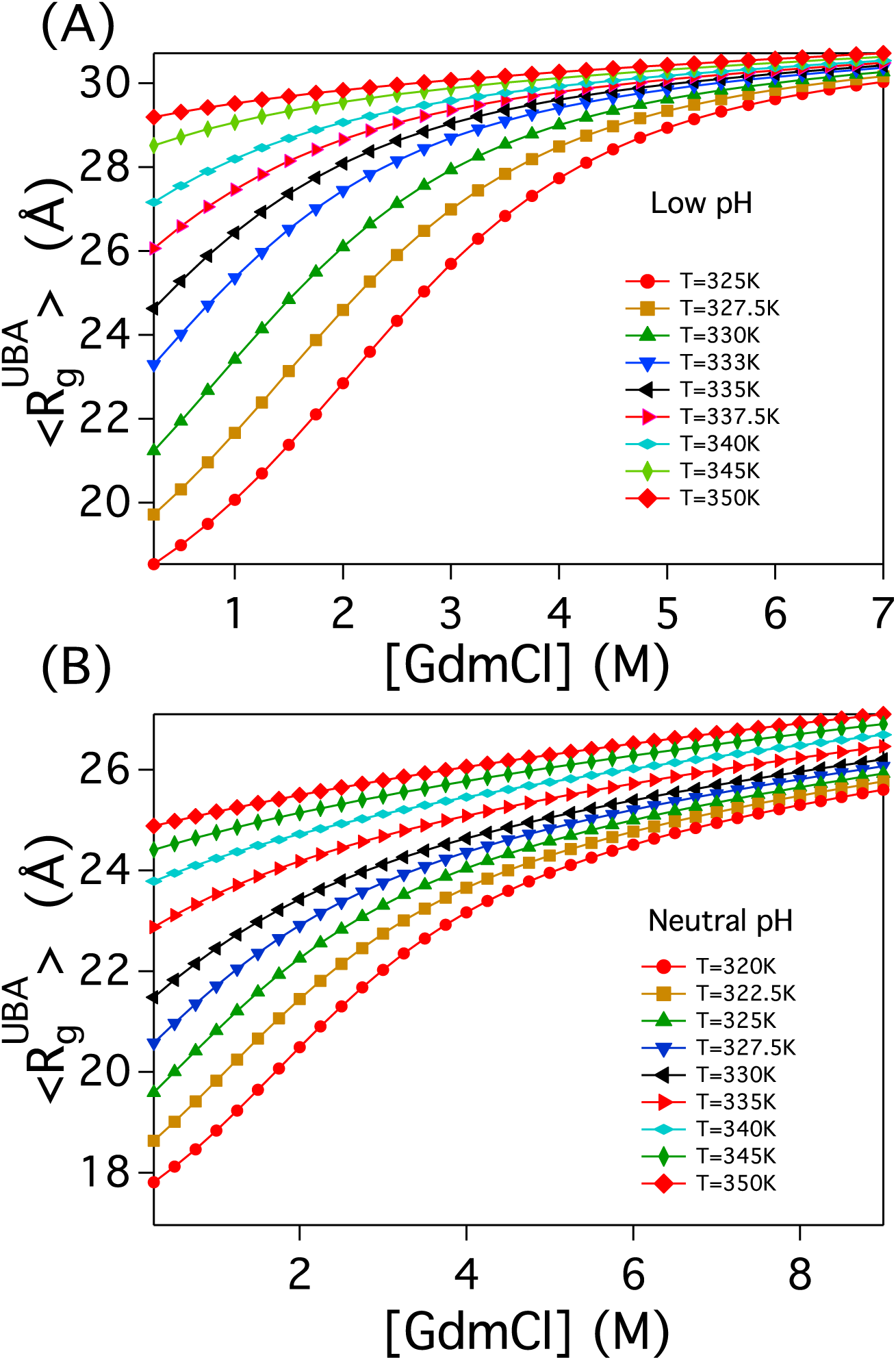
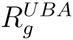 as function of [*GdmCl*] in (A) low pH and (B) neutral pH at different temperatures. The average size of the protein in the UBA basin decreases with the temperature in low [*GdmCl*].

To explore temperature effects.^70^ we calculated 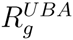 as a function of [*GdmCl*] at different values of *T*. The results in Figure 3 show that there is considerable compaction with 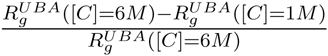 reaching nearly 30% below *T* = 333 *K* (Figure 3A). However, under experimental conditions the change is small, which would be difficult to determine accurately in typical scattering experiments. Our predictions, amenable to experimental tests, show that collapsibility can only be demonstrated by varying external conditions (for example temperature, denaturants, and force).

### Ub compaction in the the burst phase

Just as in experiments we triggered folding by decreasing [*C*] to a value below [*C*_*m*_] from a high value at which Ub is unfolded. From these dilution simulations we calculated the changes in *R*_*g*_(*t_B_|*[*C*]) during the burst-phase, *t*_*B*_, (*t*_*B*_ = 100 *µs* after initiating folding) to ascertain the extent of collapse. Each data point in Figure 4A is computed from at least 40,000 conformations from different folding trajectories. The initial unfolded protein conformations are taken from the simulations performed at [*GdmCl*] = 6 M.

**Figure 4:**
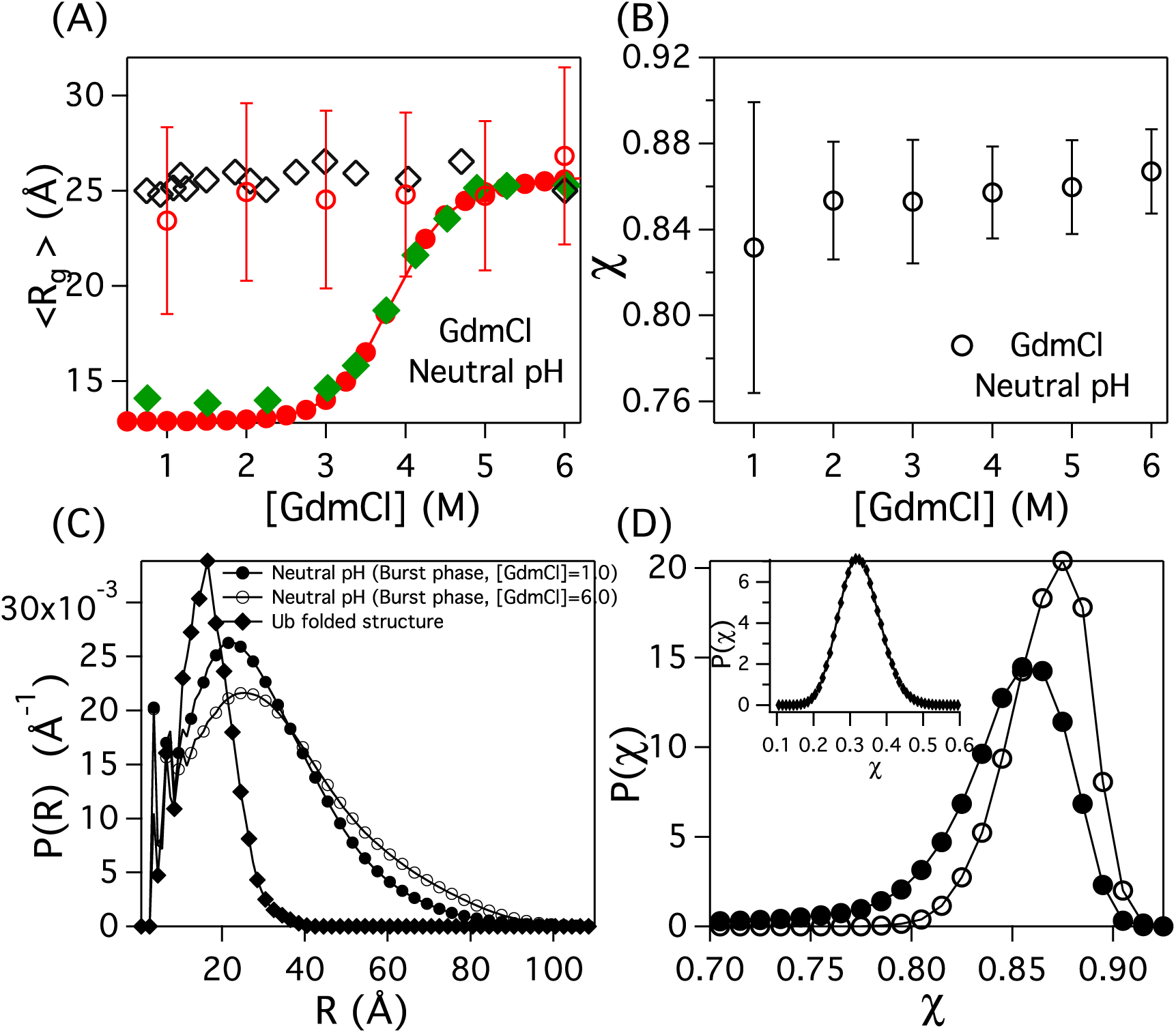
(A) Radius of gyration, *R*_*g*_, as a function of denaturant concentration. Equilibrium 〈*R*_*g*_〉 from coarse-grained simulations in neutral pH as a function of [*GdmCl*] is in red solid circles. Data in green diamonds and black empty diamonds is equilibrium and burst-phase 〈*R*_*g*_〉 as function of [*GdmCl*] from experiments^57^ at pH = 7.0. Data in empty red circles (neutral pH, *T* = 335 K, [*GdmCl*]) is 〈*R*_*g*_〉 during the burst phase of Ub folding. (B) Structural overlap function, *χ*, plotted as a function of [*C*]. The empty circle symbols in the plot represent the same conditions described in panel-(A). (C) Pair distance distribution function, *P*(*R*), plotted as a function of distance, *R*, for the Ub native structure, and during the burst phase of Ub folding. (D) Probability distribution of *χ*, *P*(*χ*). The symbols represent the same conditions described in panel-(C). Inset shows *P*(*χ*) for the folded structure in neutral pH at *T* = 335 K and [*C*] = 0 M conditions.

The data shows that during the early phase of folding, *R*_*g*_ decreases by *≈*2-3 Å in neutral pH and *T* = 335 *K* as [*GdmCl*] is diluted from 6 M to 1 M (Figure 4A). The standard deviation of *R*_*g*_ is *≈*5 Å, exceeding the observed changes in *R*_*g*_. This is because on the *t*_*B*_ time scale Ub samples conformations with widely varying *R*_*g*_ (Figure 5A). Similar behavior is observed when [*Urea*] is used as a denaturant in low pH (see Figure S3 in the SI). The experimental^57^ burst phase *R*_*g*_ data of Ub in neutral pH is in good agreement with the simulation data within the error bars. The decrease in *R*_*g*_ at low denaturant conditions is due to the partial or full folding of 2-3 trajectories within *t*_*B*_ after folding is initiated (for example trajectory in Figure 5B). During the burst phase, the structural over lap function, *χ* (defined in SI) shows that there is a small decrease in *χ* for [Urea] and [GdmCl] between 1M and 3M (Figure 4B and S3B). The standard deviation of the data between 1M and 3M is larger as only 2-3 trajectories partially or fully fold to form native contacts.

**Figure 5:**
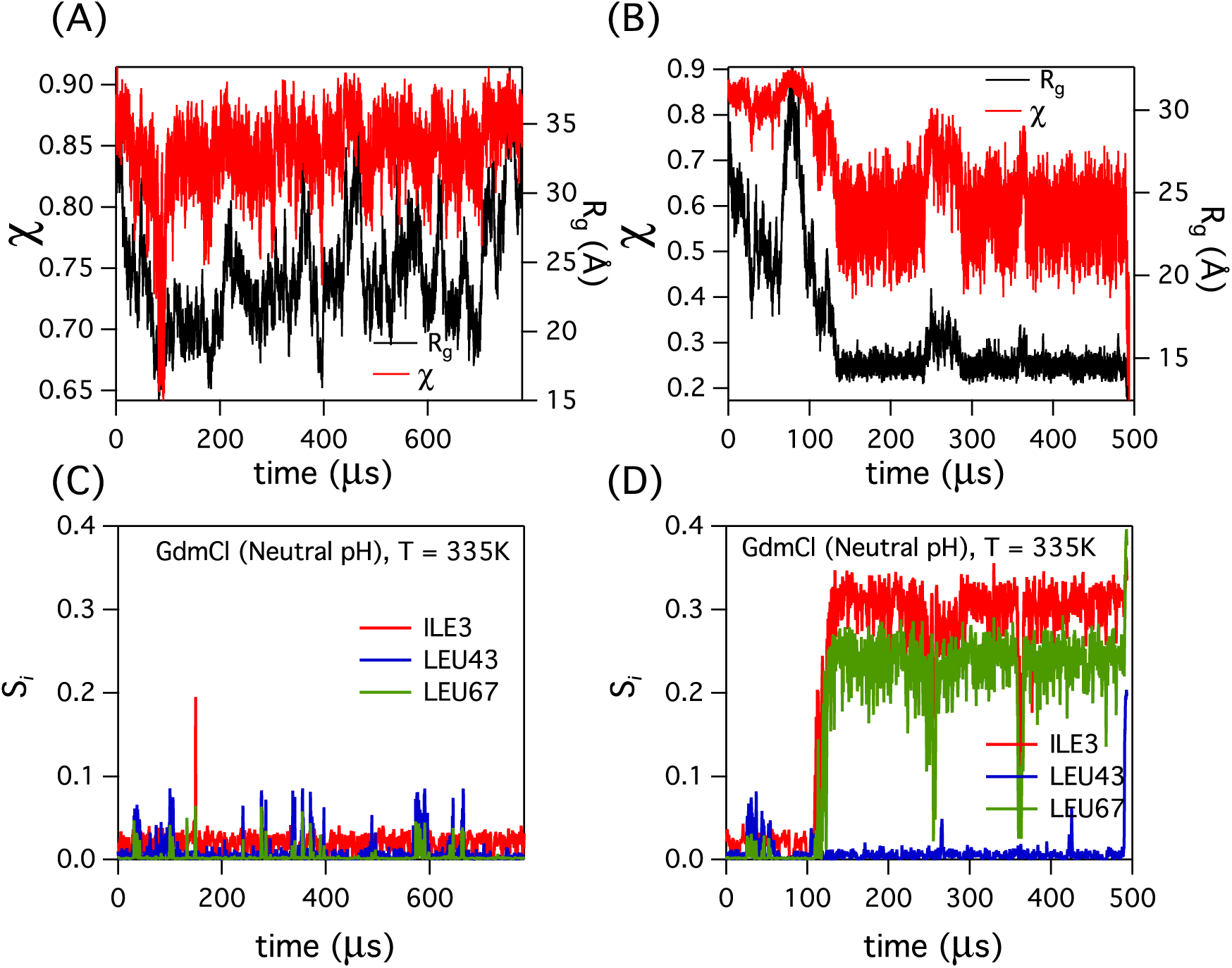
*χ* and *R*_*g*_ of a Ub folding trajectory in neutral pH is plotted as a function of time for conditions [*GdmCl*]=1.0 M and *T* = 335 *K*. (A) Trajectory where Ub does not fold in 800 *µs* and (B) trajectory where Ub folds in 500 *µs*. The contact order, *S*_*i*_, for the residues ILE3, LEU43 and LEU67 plotted as a function of time for the (C) unfolded trajectory shown in panel-(A), and (D) folded trajectory in panel-(B).

### Distance distribution functions

The pair distance distribution functions, *P*(*r*)s, the inverse Fourier transform of the scattering intensity, computed using conformations sampled at *t*_*B*_ as a function of [*GdmCl*] are in good agreement with the SAXS experiments^57^ (Figure 4C). We find that *P*(*r*) for the folded Ub spans between 0 Å *< r <* 40 Å with a peak at *≈* 16.5 Å. In contrast, *P*(*r*) in the burst phase at both low and neutral pH spans 0 Å *< r <* 100 Å with a peak at *≈* 25.5 Å showing that the protein samples expanded conformations with a large variation in *R*_*g*_. In agreement with the *P*(*r*), the probability distribution of *χ*, *P*(*χ*), also shows that native-like contacts are absent on the *t*_*B*_ time scale (Figure 4D). However, detectable compaction of Ub can be achieved by varying solvent conditions. For example, SAXS experiments^33^ performed at −20 ^*°*^C and 4 ^*°*^C in the presence of 45% ethylene Glycol show that *R*_*g*_ of the protein decreased from *≈* 26 Å to *≈* 15 Å during the burst phase of folding. Our results in Figure 3 support the conclusion reached elsewhere that environmental factors are critical in the compaction of proteins.^33^

### Compaction precedes folding

The relationship between native state formation and collapse, displayed as a two dimensional plot in *R*_*g*_ and *χ* (Figure 6), shows vividly that only upon significant decrease in *R*_*g*_ does the search for the native state begins. This finding is not only consistent with a number of rapid mixing experiments^71–73^ but also is in accord with theoretical predictions that search for the folded state occurs by sampling minimum energy compact structures.^74^ It should also be noted that even *χ ≈* 0.8, indicating that Ub is unfolded, *R*_*g*_ has decreased to about 23 Å. This shows a decrease of *≈* 8 Å (3 Å) from an average equilibrium value of 32 Å (26 Å) in low (neutral) pH (Figure 6). Thus, kinetic folding trajectories also show that compaction occurs continuously as folding conditions are changed.

**Figure 6:**
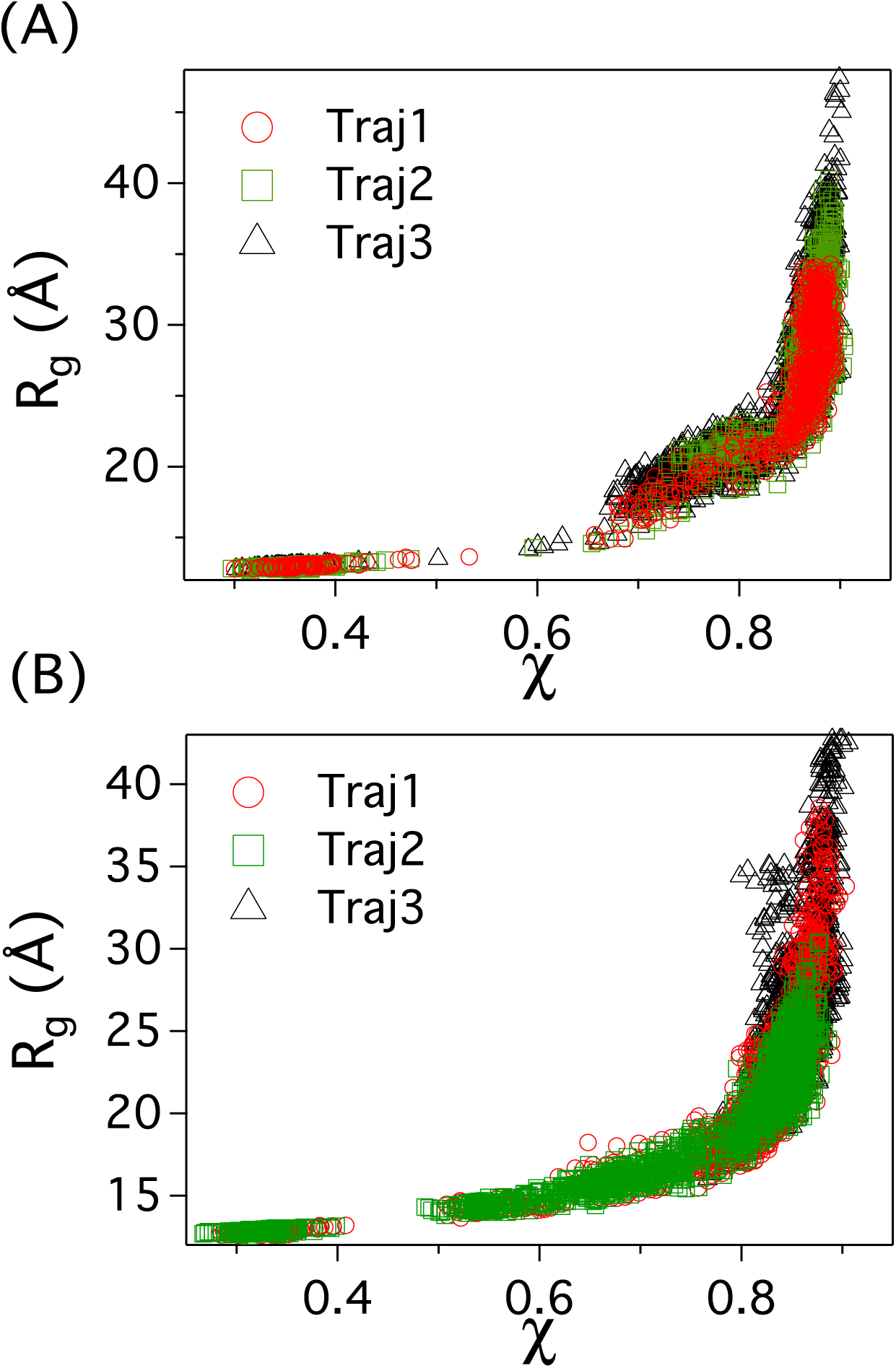
*R*_*g*_ as a function of *χ*. Three different folding trajectories are shown in red circles, green squares, and black triangles, respectively. Each data point is obtained by averaging the folding trajectory data for ≈ 0.25*µs*. (A) Low pH, T=332K, [C]=0M; (B) Neutral pH, T=335K, [C]=0M.

### Extent of compaction in the burst phase and at equilibrium is similar

The equilibrium 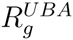 decreases continuously as the denaturation concentration decreases although the extent of compaction depends on *T* (Figure 3). Despite large errors there is detectable change in 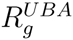 in the individual kinetic folding trajectories during the initial folding stages. Greater compaction of 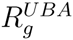 under folding conditions occurs at times that exceed the burst times, arbitrarily defined here as 100 *µs*. To illustrate the connection between compaction and formation of a minimum number of contacts we monitored the dynamics of non-local contact formation in denaturant dilution simulations.

Formation of a few such contacts is needed for compaction. On the time scale of about 100 *µs* these contacts are formed only in some folding trajectories (Figure 5). To illustrate the importance of these contacts in inducing compaction we computed the time-dependent changes in the local contact order,^64,65^ *S*_*i*_ (see methods), of the residues ILE3 (*β*_1_), LEU43 (*β*_3_) and LEU67 (*β*_5_). These residues also participate in stabilizing the hydrophobic core of the folded protein. The *S*_*i*_ for the residues ILE3, LEU43 and LEU67 shows that the long range contacts are not formed even after 800 *µs* in the folding trajectory in Figure 5A and 5C. The long range contacts, that also drive compaction, are formed only in fast folding trajectories shown in Figure 5B and 5D. More generally, although proteins that have a large fraction of non-local contacts are more collapsible (equilibrium 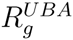 decreases more as the denaturant concentration decreases compared to proteins that are largely helical)^75^ the time scale for compaction is likely to be longer. Taken together our results firmly establish that 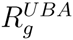 is smaller under native conditions than at high denaturant concentrations. Because the changes in 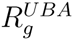 are small due to the finite size of proteins, accurate experimental measurements are needed to reach firm conclusions about the collapsibility of Ub and proteins in general.

### 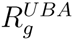 inferred from smFRET and direct simulations is qualitatively consistent

We computed the equilibrium FRET efficiency, 〈*E*〉, as a function of [*C*] in low pH by assuming that the donor and acceptor dyes are attached near the N and C termini of Ub (Figure 7A). The FRET efficiency is related to the end-to-end distance (*R*_*ee*_) probability distribution *P*(*R*_*ee*_) by,

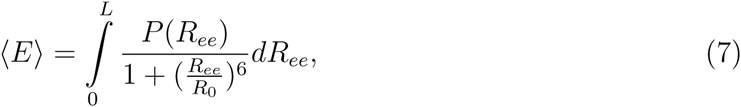

where *L*(= 285 Å) is the contour length of Ub, and *R*_0_(= 54 Å) is the Forster radius for the donor-acceptor dyes assuming the donor and acceptor dyes are AlexaFluor 488 and AlexaFluor 594, respectively.^76^ These are reasonable dyes to perform FRET experiments on Ub as *R*_*ee*_ for Ub varies between 20 Å and 130 Å, and this is approximately in the range 0.5*R*_*o*_ to 2*R*_*o*_.^77^ The FRET efficiency in the UBA, 〈*E*^*UBA*^〉, increases as the denaturant concentration, [*C*], is diluted implying that Ub becomes compact (Figure 7A and 7B).

**Figure 7:**
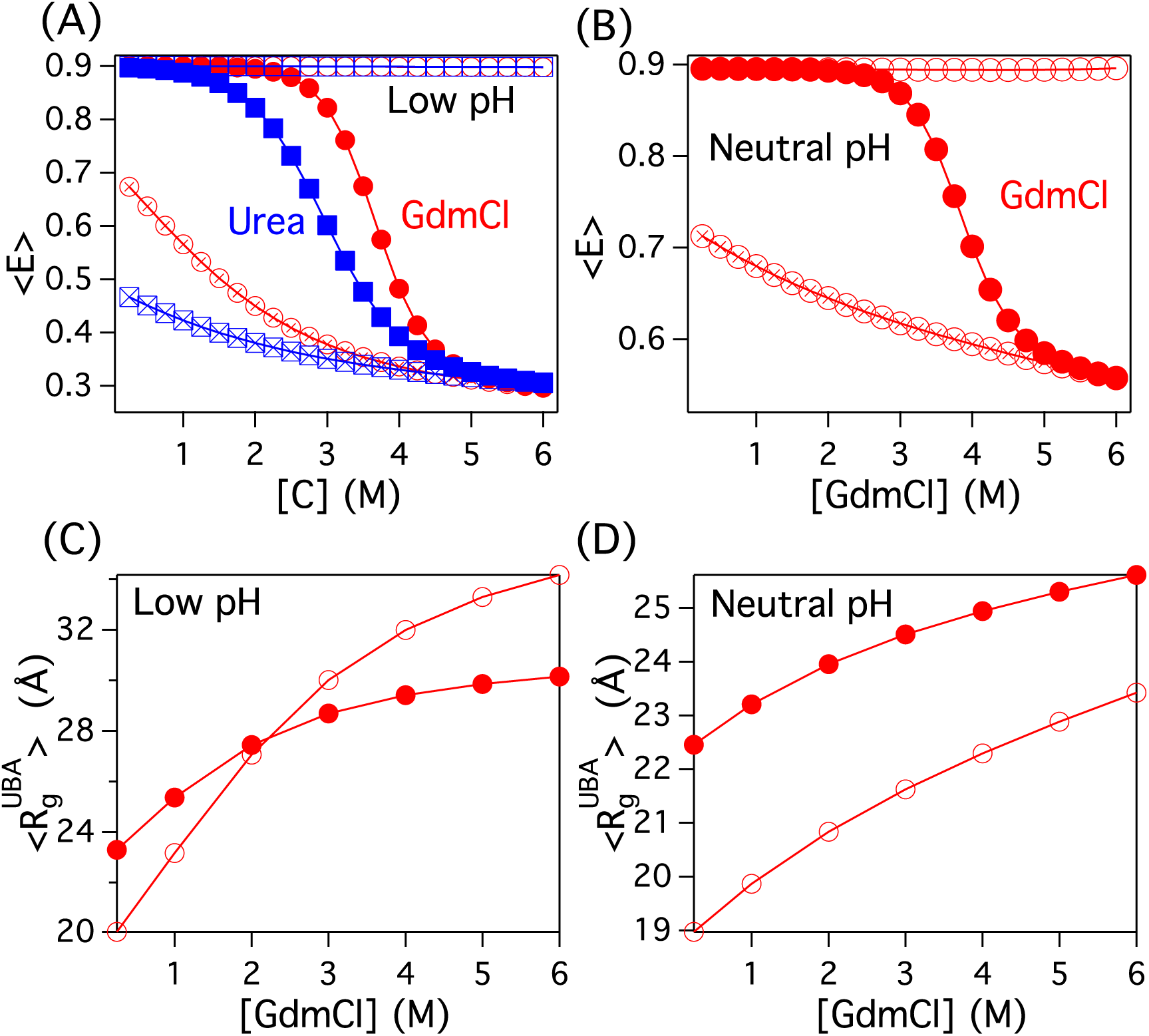
(A) Equilibrium FRET efficiency, 〈*E*〉, as a function of [*GdmCl*] and [*Urea*] in low pH is in red circles and blue squares, respectively. 〈*E*〉 for the protein conformations in the UBA and NBA basins as a function of [*GdmCl*] ([*Urea*]) is shown in cross circles (cross squares) and empty circles (empty squares), respectively. (B) 〈*E*〉, as a function of [*GdmCl*] in neutral pH. The symbols represent the same as in A. (C) Data in solid red circles is 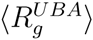 as a function of [*GdmCl*] at *T* = 333 *K* in low pH. The data is empty red circles is 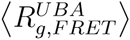 estimated from the FRET efficiency data in the UBA ensemble, 〈*E*^*UBA*^〉. (D) 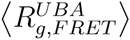 as a function of [*GdmCl*] in neutral pH at *T* = 333.25 *K*. The symbols represent the same conditions as in panel-C.

We followed the standard practice used in smFRET experiments^76,78^ to calculate 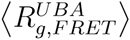 from 〈*E*〉. We assume that the unfolded state ensemble can be modeled as a Gaussian polymer with *P*(*R*_*ee*_) given by,

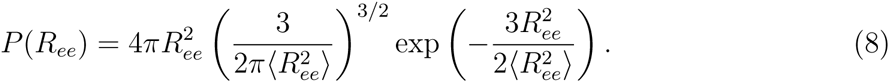

The *P*(*R*_*ee*_) in Equation 8 when used in Equation 7 yields the average end-to-end distance square, 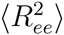. The FRET estimate for 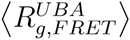 is calculated using the relation,^79^ 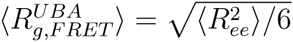.

The predicted results in low and high pH as a function of [*GdmCl*] concentration are shown in Figure 7C and 7D. Two important points are worth making: (1) In both low and neutral pH the values of 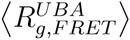 estimated from 〈*E*^*UBA*^〉 decrease as [*GdmCl*] is reduced. Thus, compaction of the unfolded proteins under folding conditions can be inferred from smFRET experiments, as we recently showed for the PDZ domain.^80^ (2) Interestingly, there is no one to one correspondence between direct calculation of 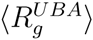 and 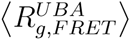. At low pH, 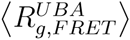 is greater than 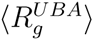 at high [*GdmCl*] whereas at a lower [*GdmCl*] the results are just the opposite. A similar finding was reported for R17 spectrin.^81^ However, at neutral pH, 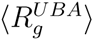 is greater than 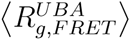 at all values of [*GdmCl*]. (3) In both low and neutral pH the extent of compaction inferred from 〈*E*〉 is greater than 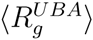, which explains the differences in the estimates of the dimensions of the unfolded state ensemble from different experimental probes.

Although the Gaussian chain end-to-end distribution gives a reasonable estimate of the *R*_*g*_ extracted from the FRET efficiency data (Figure 7C and 7D), caution should be used to make quantitative comparison with the *R*_*g*_ data estimated from SAXS. Recent FRET experiments^82^ on Ub further indicated that, although the protein shows a 2-state transition, the fluorophores lead to significant decrease in the stability of the protein pointing to the possibility that they could interact with the protein, thus effecting its stability. Furthermore for the fluorophores used in the experiment,^82^ the FRET efficiency for the protein in folded state should be *≈* 1. However, the measured value is 0.77 indicating that quenching by the residues in the vicinity of the acceptor or the orientation of the fluorophores is restricted when the protein is in the folded state. In addition, there are problems associated with the use of Gaussian chain *P*(*R*_*ee*_) to obtain absolute values of 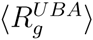 from smFRET efficiencies.^21,83–87^ Nevertheless, the qualitative conclusions obtained from smFRET experiments, which have the advantage of separating the folded and unfolded states, are valid.

### Intermediates are populated during Ub folding

In low and neutral pH and under mildly denaturing conditions (*T* = 300 K and [*GdmCl*]=1.0 M or [*Urea*]=1.0 M), Ub folding is well described by the diffusion-collision mechanism.^88^ In the early stages of folding, kinetic intermediates with folded micro domains in different parts of the protein are populated stabilized by the formation of some of the long range contacts (*β*_1_*β*_5_, *β*_3_*β*_5_ and *L*_1_*L*_2_) leading to structures I1-I4 (Figure 8) These micro domains diffuse for about 10-100 *µs* and, finally collide and coalesce to form the fully folded structure.

**Figure 8:**
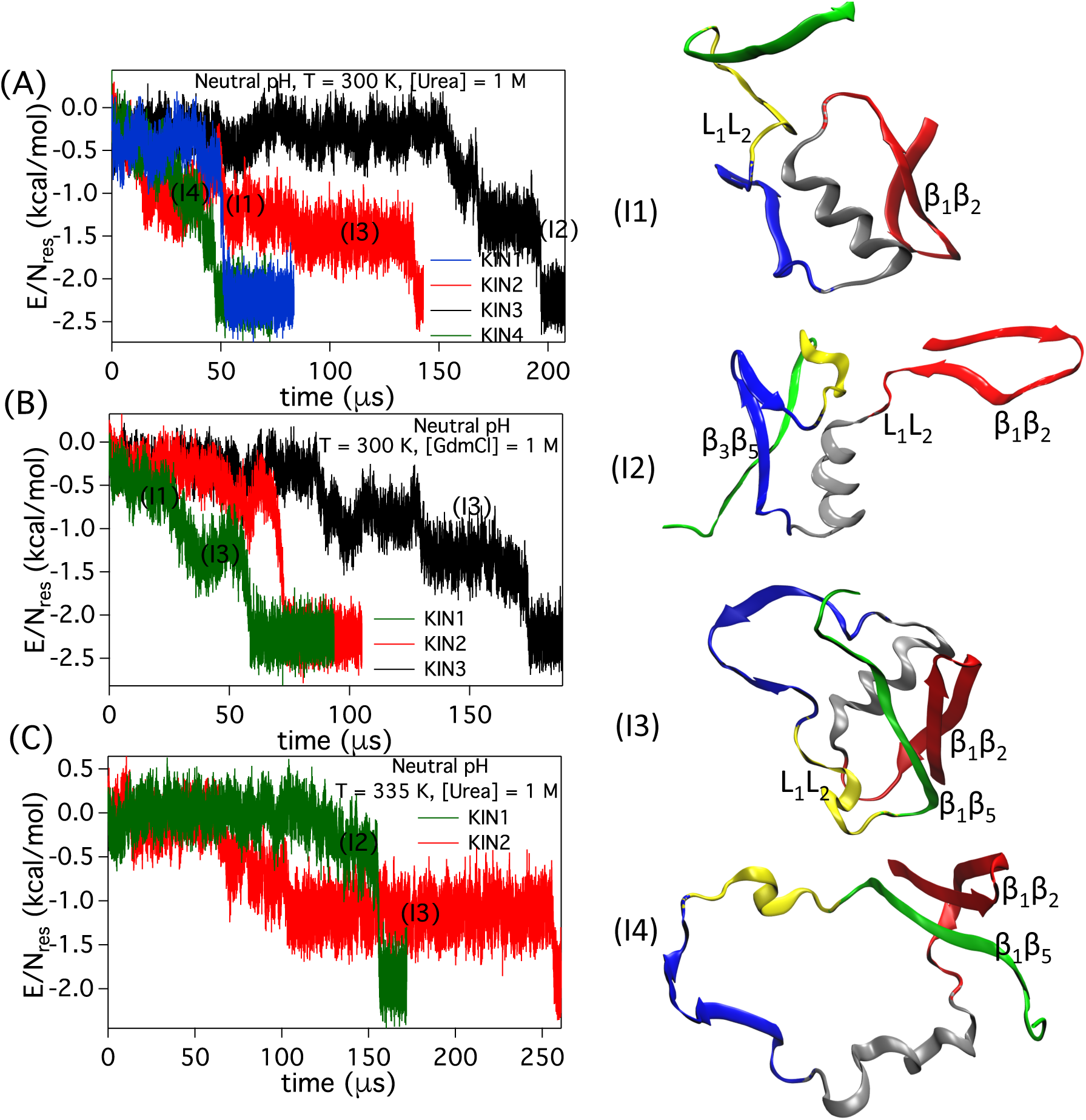
Ub folding kinetics in neutral pH. The folding pathways are inferred from change in energy per residue, *E/N_res_*, as a function of time at conditions (A) *T* = 300 *K*, [*Urea*]=1.0 M (B) *T* = 300 *K*, [*GdmCl*]=1.0 M, and (C) *T* = 335 *K*, [*Urea*]=1.0 M. Four kinetic intermediates labeled I1, I2, I3 and I4 are identified in the folding pathways. Representative structures of the kinetic intermediates are on the right.

#### Neutral pH

In highly stabilizing conditions (low *T* = 300 *K* and [*Urea*] = 1.0 M]) Ub folds through 4 different pathways (Figure 8A) suggesting that assembly occurs by the kinetic partitioning mechanism.^89^ In zero denaturant conditions at neutral pH, the first step in three of the folding pathways (KIN1-3) is the establishment of contacts between the loops *L*_1_*L*_2_ (Figure S4) resulting in intermediate I1^5^ (Figure 8). The formation of *L*_1_*L*_2_ contacts is driven primarily by the charged residues at the interface of the loops *L*_1_*L*_2_ (Figure 1A). Subsequent to the formation of *L*_1_*L*_2_ contacts, either of the long range contacts *β*_1_*β*_5_ or *β*_3_*β*_5_ form giving rise to the intermediates I2 and I3 respectively (Figure 8). In the pathway KIN4, *L*_1_*L*_2_ contacts are established within *≈* 10 *µs* after the formation of *β*_1_*β*_5_ contacts leading to the intermediate I4. In the presence of [*GdmCl*] = 1.0 M at *T* = 300 K similar structured intermediates I1-3 are observed in other folding trajectories KIN1-3 (Figure 8B and S5). The intermediates I1-3 are also observed^5^ at *T* = 300 K in zero denaturant conditions. At higher temperatures *T* = 335 K and [*Urea*] = 1.0 M, the *L*_1_*L*_2_ contacts by themselves are not very stable^5^ (Figure S6). They are formed simultaneously along with either *β*_1_*β*_5_ or *β*_3_*β*_5_ again leading to the intermediates I2 and I3 (Figure 8).

#### Low pH

In low pH and highly stabilizing conditions (*T* = 300 *K* and [*Urea*]=1.0 M or [*GdmCl*]=1.0 M), the loop contacts *L*_1_*L*_2_ form first followed by *β*_1_*β*_5_ contacts, and subsequently *β*_3_*β*_5_ contacts (Figure S7). This folding pathway gives rise to I1 and I3 (Figure 8, S8 and S9). However, at higher temperatures (*T* = 335 K and [*Urea*] or [*GdmCl*]=1.0 M) loop closure in the protein enabled by the formation of the *β*_1_*β*_5_ contacts are formed first followed by *L*_1_*L*_2_ and *β*_3_*β*_5_. This pathway populates I4 and is the dominant Ub folding pathway at the melting temperature in the absence of denaturants.^5^ This is also in agreement with the minor folding pathway inferred from the Ψ-analysis experiments^90,91^ performed at pH 7.5.

Multiple experiments pointed out the presence of kinetic intermediates in the Ub folding pathways.^24,27,33,34,92–94^ However, it is difficult to obtain a fully resolved three-dimensional structures of the intermediates from the experiments alone. There is evidence^24,27,34,92,93^ that *β*_1_*β*_2_ hairpin and the *α*_1_ helix is stable in the intermediates. In accord with this observation, our simulations show that in all the intermediates (Figure 8 there is substantial presence of the *β*_1_*β*_2_ hairpin and the *α*_1_ helix. The variation in the structures is due to the unstable *β*_3_, *β*_4_ and *β*_5_ strands, which require non-local contacts for stability. There is also experimental evidence^94^ that the small *β*_4_ strand is unstructured in the intermediates, which is also in accord with our findings (Figure 8, I1-I4). The simulations not only support the inferences made from experiments about the secondary structures in Ub kinetic intermediates but also provide additional information about the tertiary contacts in these intermediates.

## Discussion

### Unfolded Ub is compact under native conditions

Our work shows that the unfolded 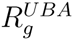 decreases continuously as [*C*] changes from a high to a low value (Figure 3). However, the extent of compaction depends on temperature. For example, 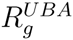 changes only by *≈* 3 Å at *T* = 335 K, a condition chosen to obtain the experimentally measured stability. The change in 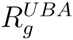 is only *≈* 10% reduction relative to [*GdmCl*] = 7M. By lowering *T* to 325 K, a 30% reduction in 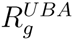 is predicted when [*GdmCl*] decreases from 7M to 1M. These findings show that 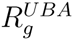 always decreases as the denaturation concentration is lowered but the extent of compaction depends on the temperature.

An argument used in the previous studies claiming the absence of collapse in Ub^57^ is that the equilibrium *R*_*g*_ and the burst phase *R*_*g*_ coincide. This is indeed the case in our simulations as well. The magnitude of change in *R*_*g*_ during the burst phase in our simulations is just as small as under equilibrium but is computable. Given that that there is only a modest decreases in 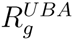 at *T* = 335 K there ought to have been a more precise analysis of the SAXS data with error estimates to rule out the claim that Ub does not collapse.^57^ Absent such an analysis we believe that the conclusions reached elsewhere are at best tenuous.^57^ Our findings suggest that additional experiments are needed to assess the propensity of Ub to collapse.

### Finite size effects and Experimental issues

Using theoretical arguments we showed^95,96^ that finite size of proteins plays a major role in restricting the magnitude of changes in the dimensions of unfolded proteins under folding conditions. In particular, theory and simulations have shown that 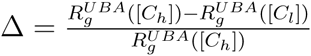 is typically small under conditions explored in experiments. For Ub, we find that ∆ *≈* 0.13 (see Table 1). Recent experiments^81^ on R17 spectrum show that ∆ *≈* 0.17 is also small. Indeed, several proteins for which data or reliable simulation results are available the values of ∆ are not large (Table 1). Thus, experiments that can distinguish the structural characteristics of the unfolded states from the folded conformations at [*C*] *<* [*C*_*m*_] are needed to establish the extent of collapse. Clearly this is more easily realized in smFRET experiments. However, the current method of extracting 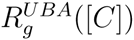 using polymer model for *P*(*R*_*ee*_) could overestimate (underestimate) the size of the unfolded states of proteins at high (low) values of [*C*], thus exaggerating the extent of compaction. In addition, the attachment of dyes could also compromise the stability of the protein as suggested in a recent study on Ub.^82^ Nevertheless, the conclusions inferred from smFRET experiments appear to be qualitatively robust, and are in line with SAXS measurements.^81^

**Table 1:** Relative changes in the radius of gyration of various proteins in the UBA ensemble 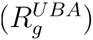 between high and low denaturant concentrations

| Protein | $R_g^{UBA}([C_h])^d$<br>([GdmCl] = 7 M) | $R_g^{UBA}([C_l])^e$<br>([GdmCl] = 1M) | $\Delta = \frac{R_g^{UBA}([C_h]) - R_g^{UBA}([C_l])}{R_g^{UBA}([C_h])}$ |
| --- | --- | --- | --- |
| Protein L<sup>21</sup> ($N_{res}=64$) | 26.3 Å | 22.5 Å ([C]=0.25 M) | 14.4% |
| Monellin ($N_{res}=96$; Neutral pH)<sup>a</sup> | 27.8 Å | 25.5 Å ([C]=0.25 M) | 8.3% |
| PDZ2 Domain<sup>80</sup> ($N_{res}=94$)<sup>b</sup> | 32.2 Å | 29.8 Å ([C]=0.25 M) | 7.5% |
| Ubiquitin ($N_{res}=76$; Neutral pH) | 25.9 Å | 22.5 Å ([C]=0.25 M) | 13.1% |
| Ubiquitin ($N_{res}=76$; Low pH) | 30.4 Å | 23.3 Å ([C]=0.25 M) | 23.4% |
| SH3<sup>19</sup> ($N_{res}=56$) | 23.7 Å | 20.3 Å ([C]=0.2 M) | 14.3% |
| Cold Shock<sup>16</sup> ($N_{res}=70$) | 26.4 Å | 17.8 Å ([C]=1 M) | 32.6% |
| ACTR<sup>81</sup> ($N_{res}=73$)<sup>c</sup> | 29.7 Å | 24.7 Å ([C]= 0.3 M) | 16.8% |
| R17d<sup>81</sup> ($N_{res}=116$)<sup>c</sup> | 40.3 Å | 33.2 Å ([C]= 0.6 M) | 17.6% |

<sup>a</sup> Unpublished data
<sup>b</sup> Denaturant used is Urea
<sup>c</sup> Data from experiments
<sup>d</sup> $R_g^{UBA}([C_h])$ denotes $R_g$ of the protein in the UBA ensemble in high denaturant concentrations
<sup>e</sup> $R_g^{UBA}([C_l])$ denotes $R_g$ of the protein in the UBA ensemble in low denaturant concentrations

Table 1 shows that there is little correlation between ∆ and the length of the proteins, which have been studied experimentally to quantitatively estimate the extent of compaction. Not only is the variation in length small so are the changes in 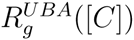 as [*C*] is decreased from a high to a low value less than *C*_*m*_. The extent of relative compaction (∆) varies from about 8% to 33%. The maximum change occurs in cold shock protein with a *β*-sheet architecture in the folded state. This observation accords well with recent theoretical predictions^75^ establishing that collapsibility in proteins with *β*-sheet structure is greater than in *α*-helical proteins.

### Fate of Ub in the early stages of folding

The folding trajectory in Figure 5 shows that on the time scale of about 100 *µ*s there is a reduction in 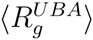 without accumulation of native-like structure as measured by *χ*. The extent of reduction coincides with the estimates based on results obtained at equilibrium. To set this observation in a broader context, it is useful to consider the relevant time scales when refolding is initiated by diluting the denaturant concentration. Broadly, we need to consider the interplay of three time scales. These are the folding time (*τ_F_*), the collapse time (*τ_c_*), and the time for forming the first tertiary contacts (*τ_tc_*) required to decrease 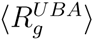. The folding trajectory (Figure 8 and S4) shows that when *τ_tc_ ≈ τ_c_ ≈* 50 *µ*s the radius of gyration decreases while the energy per residue is at least four times greater than the value in the folded state. Thus, compaction occurs without any sign of folding. In addition, the [*R_g_, χ*] plot shows that on the time scale *τ*_*c*_ the value of *χ* is far greater than that in the folded state. Taken together, these results imply that Ub is compact on the burst time scale without being in the NBA.

Our predictions for Ub are consistent with several previous experiments on other proteins. (1) In a recent study on Monellin, Udgaonkar has shown^97^ that on the *τ_c_ ≈* 37 *µ*s (the folding time is considerably longer) the polypeptide chain contracted without much structure. These compact structures have been suggested to be minimum energy compact structures (MECS) predicted theoretically.^74^ (2) Using site specific hydrogen-deuterium (H/D) in horse Cytochrome-c (Cyt-c) Roder and coworkers^98^ showed that on a time scale of *τ_c_ ≈* 140 *µ*s (*< τ_F_*, which exceeds several *ms*) the protein collapses with establishment of native-like local contacts. This work also showed that Cyt-c compaction is unlikely to be due to the presence of the heme group. Rather, it is an intrinsic property of the protein. Interestingly, our simulations on Ub show that specific contacts are necessary for a reduction in 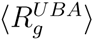. Thus, it is likely that a universal mechanism of compaction of polypeptide chain involves formation of few native contacts. (3) In order to rationalize the interpretation based on SAXS experiments that collapse is absent in globular proteins it has been asserted that the exceptions hold only for proteins with disulfide bonds in the native state or proteins with prosthetic groups undergo collapse. This argument contradicts the recent discovery, using single molecule pulling experiments,^99^ showing that disulfide bond forms only after compaction of the polypeptide chain. The experiments are supported by theoretical predictions of folding of BPTI,^100^ which have been further substantiated using simulations.^101^ Thus, both experiments and computations established that prior to the formation of the first disulfide bond there is reduction in the dimensions of the protein and not the other way around. The overwhelming evidence suggests that unfolded states of proteins are compact under native conditions with the extent of compaction being modest (Table 1).

### Kinetic intermediates in refolding of Ub

It has remained controversial if Ub folds by populating discernible intermediates, whose formation of intermediates does depend on a number of factors. For example, Khorasanizadeh^35^ showed that under stabilizing conditions the refolding kinetic data on Ub can only be explained by a three state model. Our simulations suggest that the kinetics could be even more complex. Folding occurs by parallel routes described by the KPM in which a fraction of molecules folds in a two-state manner whereas others reach the native state by populating distinct intermediates. These results suggest that only by varying the stability conditions and using high temporal resolution experiments can the complexity of Ub folding be fully elucidated.

## Conclusions

We performed molecular dynamics simulations using the coarse-grained SOP-SC model and molecular transfer model of Ubiquitin folding in Guanidinium Chloride and Urea solutions to address two questions of fundamental importance in protein folding. One of them is concerned with the size of the unfolded states under native conditions. We showed that under all conditions Ub does become compact as the denaturant concentration is decreased with the extent of compaction being dependent on stability of the folded state. There is complete consistency between equilibrium and burst phase values of the radius of gyration of the unfolded states. Interestingly, the mechanism of Ub collapse is due to the formation of specific contacts in the unfolded state. In other words, the structure of the unfolded state under folding conditions is more compact than at high denaturant concentration. These and related experimental and computational studies show that the propensity for globular proteins to collapse is universal.

Our simulations, which are in quantitative agreement with experiments,^36,39^ also addresses the second question, namely, are there intermediates in Ub folding? The answer to this question is in the affirmative. The kinetic intermediates observed during the folding pathways are identical to the ones observed in the temperature induced folding pathways in the absence of denaturants. The structures of the kinetic intermediates are determined by a combination of the long range contacts between the secondary structural elements *L*_1_*L*_2_, *β*_1_*β*_5_ and *β*_3_*β*_5_. Thus, folding of Ub is more complicated than previously thought but is well described by the kinetic partitioning mechanism.^89^ These predictions can be validated using experiments with high temporal resolution. Finally, the combination of coarse-grained simulations and the use of MTM to account for denaturants is a major advance in examining folding of a variety of proteins in quantitative detail.

## Acknowledgement

GR acknowledges startup grant from Indian Institute of Science-Bangalore, and funding from Nano mission, Department of Science and Technology, India. DT acknowledges a grant from the National Science Foundation through grant CHE 16-36424 and the Welch Chair at the University of Texas at Austin. Some of the computations are performed by C-DAC’s PARAM Yuva-II. A portion of this research used resources of the National Energy Research Scientific Computing Center, a DOE Office of Science User Facility supported by the Office of Science of the U.S. Department of Energy under Contract No. DE-AC02-05CH11231.

## Supporting Information Available

Description of the simulation methods; Table S1; Figures S1-S9. This material is available free of charge via the Internet at http://pubs.acs.org/.

---

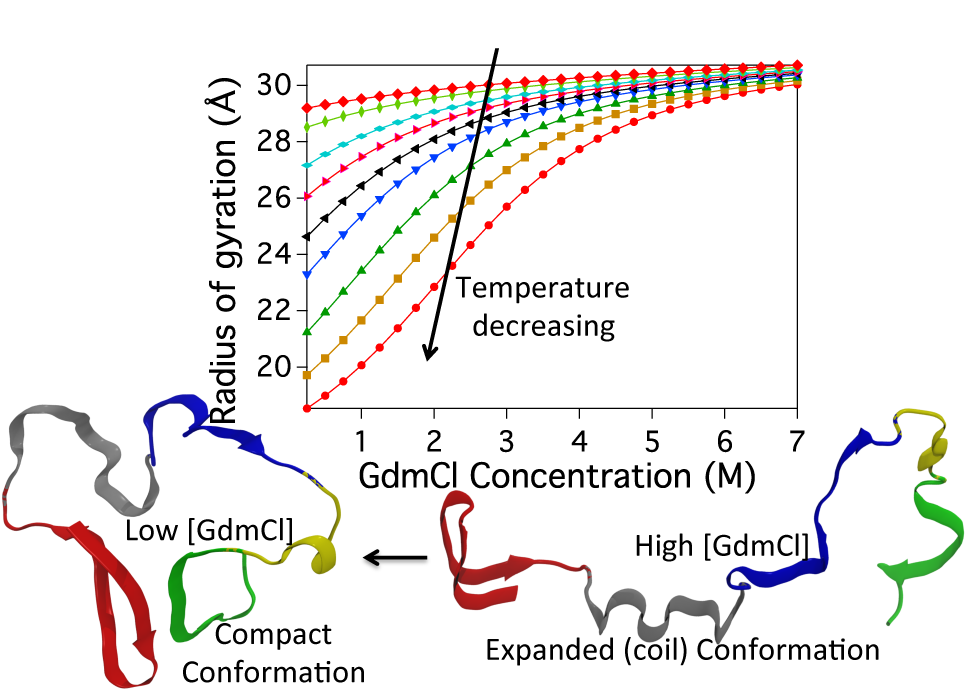
Table of Contents (TOC) figure

